# Local and Global Measures of Information Storage for the Assessment of Heartbeat-Evoked Cortical Responses

**DOI:** 10.1101/2023.06.07.544078

**Authors:** Chiara Barà, Andrea Zaccaro, Yuri Antonacci, Matteo Dalla Riva, Alessandro Busacca, Francesca Ferri, Luca Faes, Riccardo Pernice

## Abstract

**Objective:** Brain-heart interactions involve bidirectional effects produced by bottom-up input at each heartbeat, and top-down neural regulatory responses of the brain. While the cortical processing of the heartbeat is usually investigated through the analysis of the Heartbeat Evoked Potential, in this study we propose an alternative approach based on the variability in the predictability of the brain dynamics induced by the heartbeat.

**Methods:** In a group of eighteen subjects in whom simultaneous recording of the electroencephalogram (EEG) and electrocardiogram was performed in a resting-state, we analyzed the temporal profile of the local Information Storage (IS) to detect changes in the regularity of EEG signals in time windows associated with different phases of the cardiac cycle at rest.

**Results:** The average values of the local IS were significantly higher in the parieto-occipital areas of the scalp, suggesting an activation of the Default Mode Network, regardless of the cardiac cycle phase. In contrast, the variability of the local IS showed marked differences across the cardiac cycle phases.

**Conclusion:** Our results suggest that cardiac activity influences the predictive information of EEG dynamics differently in the various phases of the cardiac cycle.

**Significance:** The variability of local IS measures can represent a useful index to identify spatio-temporal dynamics within the neurocardiac system, which generally remain overlooked by the more widely employed global measures.

## 1 Introduction

Complexity and dynamics are two of the main properties of the processes measured at the output of realworld systems including human organisms. These properties arise from the natural evolution over time of the several intertwined physiological systems that compose the human organism [1, 2]. In fact, the continuous changes in the mutual activity of several control mechanisms across different physiological states and pathological conditions typically result in non-linear, time-varying and multiscale dynamic behaviors that are difficult to understand by looking separately at each organ alone [3, 4, 5, 6, 7, 8, 9, 10, 11, 12, 13, 14, 15]. This evidence has promoted studies, framed in the emerging field of Network Physiology [5, 16, 3], addressing the human organism as a network of horizontally integrated systems, each having an internal/vertical organization. To investigate the complex dynamics emerging from physiological interactions, tools developed for the analysis of complex systems are typically adopted (e.g., fractal dimension [17], Lyapunov exponents [18], Lempel-Ziv complexity [19]). Among these tools, entropy metrics are gaining increasing popularity thanks to their generality and flexibility, and to their applicability to short and noisy realizations of physiological stochastic processes [20]. These measures allow to quantify the information content of a dynamic system starting from the probability distribution of the variables that describe the temporal evolution of its states. One important measure providing information about the complex dynamics generated by a stochastic process is the Information Storage (IS), defined as the information content of the current state of the process that can be used to predict its future. Thus, IS allows to quantify the regularity and predictability of a dynamic process [21, 22, 23, 24]. Together with measures quantifying the information transfer between coupled processes, the IS constitutes a basic element of computation to assess the processing of information within networks of multiple interacting systems [22, 24]. These measures have been widely employed to investigate several physiological mechanisms, such as those underlying cardiovascular [8, 21, 25, 26], cardiorespiratory [9, 24, 27], and neural [10, 28] dynamics.

Measures like the IS allow to extract information about the dynamics of the observed process analyzed as a whole [29], thus representing “global” indexes which address the entire temporal evolution of the process. This approach is often followed in the analysis of physiological time series as it provides a compact description of the process based on a single IS measure. However, the global approach is limited in the fact that it does not allow to obtain time-resolved information about the process dynamics. This limitation can be addressed through the definition of time-varying information measures which, relaxing the assumption of stationarity, evaluate the dynamics at each specific time step accounting for the data collected in a preceding short temporal window [11]. However, time-varying measures have the disadvantages to use a limited portion of the data for the estimation, and to depend strongly on the tradeoff between temporal resolution and ability to detect the local temporal properties of the analyzed process. An alternative, theoretically well-grounded approach is the use of local measures of information dynamics [24]. This approach defines how information is stored, transferred and modified at each moment in time in a multivariate stochastic process [22, 30, 31]. In particular, measures of local IS and local information transfer are computed from the joint and marginal probability distributions of the variables mapping the present and past states of the analyzed processes computed at any specific time step, such that their statistical average yields the global measures of IS and of transfer entropy. Local measures computed from single realizations of physiological processes analyzed under the assumption of stationarity have been widely applied to characterize specific alterations of brain activity in cats during visual stimulation [23] or in humans with autism spectrum disorders [12], to detect phase-amplitude coupling in cortical local field activity [32], or to analyze time-resolved properties of cardiorespiratory interactions during sleep apneas and of the cortical information flow during focal epilepsy [33].

Physiologically, brain-heart interactions are often studied by investigating the regulatory efferent activity of the Autonomic Nervous System (ANS) on the cardiac rhythm, given that the descending control of the brain towards the heart determines heart rate variability [34, 35, 36]. Nevertheless, the communication between brain and heart is bidirectional [6, 37, 38], since an ascending control is also present, whereby neural activity is influenced by cardiac activity [38]. In fact, several studies suggest how heart timing enables the optimization of numerous neural processes related to homeostatic and allostatic regulation [39]. In addition to the ANS and its sympathetic and parasympathetic branches, other mechanisms of communication between heart and brain concern baroceptors located in the aortic arch and the carotid arteries [40], cardiac neurons present in the heart wall, cutaneous receptors in the skin [40, 41] and the Intrinsic Cardiac Nervous System (ICNS) [42]. Indeed, a brain-like structure, including ganglia, neurotransmitters, proteins, and cells, is located inside the heart and is able to integrate signals from the ANS and all sensory systems, collecting information that concerns the cardiovascular activity. Signals integrated by the ICNS reach the brain and the nucleus tractus solitarii (NTS) via the vagus nerve and the spinal cord, ending in specific cortical regions [34, 43]. In fact, functional imaging techniques have allowed so far to identify targets of visceral signals in the ventromedial prefrontal cortex (vmPFC), in the insular cortex and in the motor cingulate region, corresponding to the anterior cingulate cortex (ACC) [34, 44, 45, 46]. These cortical regions partially coincide with the two brain structures mainly involved in brain-heart interactions, i.e., the Central Autonomic Network (CAN) and the Default Mode Network (DMN) [39]. The CAN represents a group of brainstem and cortex regions involved in cognitive and autonomic regulatory mechanisms, which result in interactions with the ANS through feed-forward and feedback loops, shaping bidirectional communication between the brain and the body [47]. The DMN constitutes a group of brain regions characterized by higher intrinsic activity during a resting-state condition rather than during the execution of a task [48, 49, 50]. Within the DMN, the vmPFC receives sensory information from both the internal body (interoceptive) and the external environment (exteroceptive), and transmits them to hypothalamus and amygdala. Hence, this cortical region plays a key role in the link between sensory and visceral-motor function, as well as in behavioral control [51]. This suggests that the study of the cortical processing of the heartbeat may have strong implications for our understanding of perceptual, cognitive and emotional processes.

The present study introduces an approach for the computation of local IS measures tailored for the analysis of the effects of the heartbeat on cortical dynamics measured from the scalp electroencephalogram (EEG). We first employ a strategy for estimating the local IS from single-trial EEG recordings, based either on the identification of a linear autoregressive (AR) model, or on the computation of the local information content through a non-parametric nearest neighbor technique. Then, to analyze the regularity of the neural activity timed with the heartbeat, we investigate the time series of local IS by computing their mean and variability within specific temporal windows synchronized with each heartbeat detected from the electrocardiogram (ECG). Our approach investigates brain-heart interactions from a new perspective, focused on the impact that the heartbeat has on the local EEG dynamics during different phases of the cardiac cycle. Moreover, the utilization of multichannel EEG leads us to build maps of brain-heart interaction which highlight regions of the scalp that are most interested by changes of the local IS patterns evoked by the heartbeat.

## 2 Material and methods

### 2.1 Information-theoretic preliminearies

The information-theoretic analysis of time series is grounded on basic concepts of information theory that are recalled in the following. A central measure of information theory is the entropy of a random variable [20], which measures, in a statistical sense, the average information content of the variable. Specifically, given a (possibly vector) random variable *V*, the Shannon entropy is defined as *H*(*V*) = *−*𝔼[log *p*(*v*)], where *p*(*v*) is the probability density function (PDF) of *V* measured for the outcome *v*, and 𝔼[*·*] is the expectation operator taking the statistical average over all possible values *v* taken by *V*. A more specific quantity is the information content of the single outcome *v*, which is defined as *h*(*v*) = *−* log *p*(*v*). This measure allows a *local* analysis of the information content of a random variable, i.e., an analysis focused on a specific outcome of the variable, while the Shannon entropy can be interpreted as a global measure, as it corresponds to the average information content, i.e., *H*(*V*) = 𝔼[*h*(*v*)].

The considerations above can be extended to any information theoretic measure, so as to interpret a global measure as the statistical average of its local counterpart. In particular, the conditional entropy (CE) of *V* given another variable W quantifies the residual information about *V* when *W* is known as the average uncertainty that remains about *V* when the outcomes of *W* are assigned: *H*(*V* |*W*) = 𝔼[*h*(*v*|*w*)], where *h*(*v*|*w*) = *−* log *p*(*v*|*w*) is the local CE. Similarly, the mutual information (MI) quantifies the information shared between *V* and *W* as the average uncertainty about one variable that is resolved by knowing the other: *I*(*V*; *W*) = 𝔼[*i*(*v*; *w*)], where 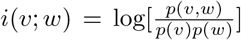 is the local MI. Note that entropy, CE and MI are linked to each other in both their local and global formulations, i.e., *h*(*v*) = *i*(*v*; *w*)+ *h*(*v*|*w*) and *H*(*V*) = *I*(*V*; *W*)+*H*(*V* |*W*). Note also that, while the MI is always non-negative, the local MI can take both positive and negative values; in the latter case, learning the outcome *w* for the variable *W* is interpreted as misinformative about the specific outcome *v* of the variable *V* [22]. In this study, the natural logarithm is used to compute information-theoretic measures, which are thus expressed in nats.

### 2.2 Global and Local Information Storage

Let us consider a dynamic system 𝒳, and assume that the evolution of the system over time is described by the stochastic process *X* = {*X*_*n*_}, *n* ∈ ℤ. Considering the temporal sampling of the process, we assume the scalar variable *X*_*n*_ and the vector variable 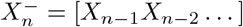 as representative of the present state and of the past states of the process, respectively. Then, a measure of the regularity of the process is the so-called information storage (IS) defined as [1, 22]:

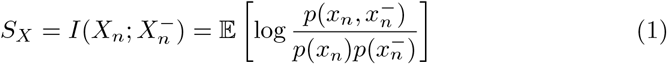

where *x*_*n*_ and 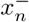 are realizations of *X*_*n*_ and 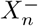. The IS quantifies the predictability of *X* intended as the average uncertainty about the current state of the process that could be resolved from the knowledge of its past states. Note that the definition of IS is provided in (1) for a stationary process *X*, for which the MI is independent on the time step *n*.

Being defined as a MI, the IS can be expressed as the statistical average of a local MI, i.e., 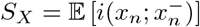, which evidences the so-called local IS [22, 30]:

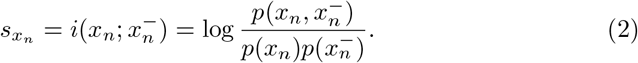

The local IS 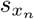 represents the amount of storage information used by the process in the specific time instant *n*. While the global measure of IS always assumes positive values and is bounded above by the entropy of the current state of the system, its local correspondent is not bounded and could assume negative values [22, 30]. In particular, positive values of 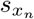 occur when knowing the past state 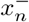 increases the probability of observing the present state *x*_*n*_, while negative values of 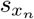 are measured when the past of the process is misinformative about the current state and thus reduces its predictability.

In practical applications, the local and global IS measures are computed for stationary processes under the hypothesis of ergodicity starting from a single process realization available in the form of the finite-length time series *x* = {*x*_1_, …, *x*_*N*_}. Moreover, since infinite-length histories cannot be considered, the assumption that *X* is a Markov process with finite memory *q* is typically made, so as to approximate the past history of the process with the *q*-dimensional variable 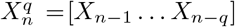. In such a case, the global IS becomes 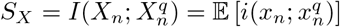, where the local IS 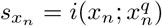 is computed approximating the past history relevant to the sample *x*_*n*_ with the vector 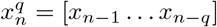. Under these assumptions, the local IS measure is estimated as:

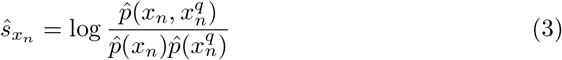

where an appropriate estimate 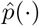 of the joint and marginal probabilities is used, and the global IS estimate is then obtained as the temporal average of the local IS

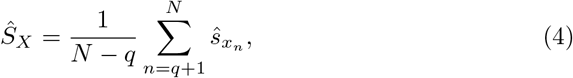

where the sum is extended to all patterns 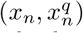 that can be derived from the time series *x* of length *N*. Depending on the hypothesis made about the data distribution, several types of estimators can be used. In this work, a parametric and a model-free approach are exploited, respectively the linear and the k-nearest neighbor estimators, which are described in the following subsections.

### 2.3 Linear Parametric Estimation

The parametric approach assumes a specific shape of the probability density function, making it possible to estimate information measures knowing the parameters of such distribution [20]. In particular, the assumption of a joint Gaussian distribution for the variables forming the present and past states of the observed process is very useful as it is strictly related with the linear parametric representation of the process [52]. The PDF of a *d*-dimensional Gaussian random variable *V* = [*V*_1_ … *V*_*d*_] with mean *µ*_*V*_ = 𝔼[*V*] and covariance matrix **Σ**_*V*_ = 𝔼 [(*V − µ*_*V*_)^⊤^(*V − µ*_*V*_)] is given by:

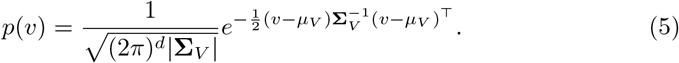

This formulation can be exploited, for the zero-mean stationary process *X*, to write the PDF of the variables *X*_*n*_ and 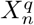 involved in the definition of the IS, as well as their joint PDF, as:

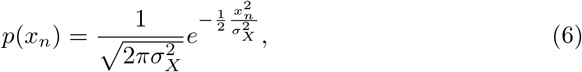

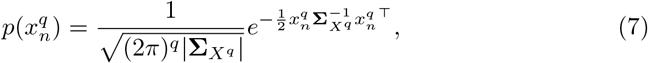

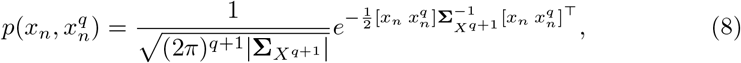

where 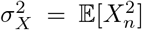 is the variance of 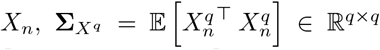 is the covariance matrix of 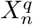 and 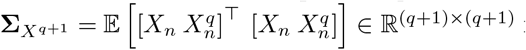 is the joint covariance matrix of *X*_*n*_ and 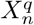. Then, an explicit formulation of the local IS for Gaussian processes can be obtained substituting these PDFs into the definition:

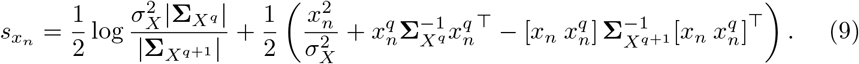

It is possible to show [33] that the first term in (9) corresponds to the global IS, i.e.,

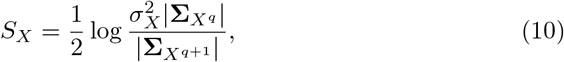

and that the second term in (9), which changes at any time step *n*, has zero average.

Given (9) and (10), we can see that the computation of the local and global IS amounts to computing the relevant covariance and cross-covariances matrices between the present and the past variables of the process, i.e., 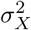, **Σ**_*X*_*q*, and **Σ**_*X*_*q*+1. These matrices can be derived from the autocovariance structure of the linear autoregressive (AR) representation of the process *X* [27]:

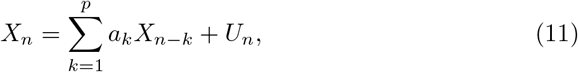

where *p* is the AR model order, *A* = [*a*_1_ … *a*_*p*_] ∈ ℝ^1*×p*^ is the coefficient vector and *U* is a white Gaussian noise process with zero mean and variance 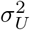. The autocovariance of the process (11) is related to the AR parameters via the Yule-Walker equations [53]:

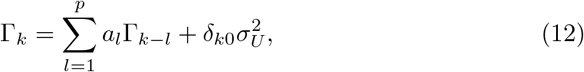

where Γ_*k*_ = 𝔼[*X*_*n*_ *X*_*n−k*_] represents the autocovariance of the process defined at each lag *k ≥* 0, *δ*_*k*0_ is the Kronecher product and 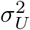 the variance of *U*_*n*_. In order to determine the autocovariance of the process for each lag *k*, the AR model can be written compactly [27] as 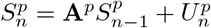 where:

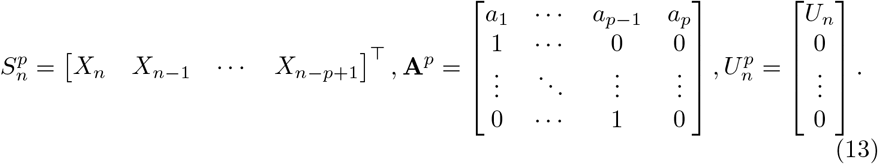

Specifically, the covariance matrix of 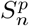 can be expressed as a discrete-time Lyapunov equation:

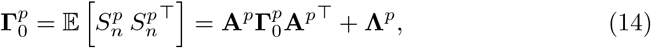

where 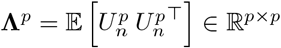 is the covariance matrix of 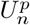 depending only on 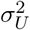. Solving (14) for 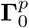 allows to obtain autocovariance values for lag between 0 and *p −* 1, **Γ**_0_ … **Γ**_*p−*1_; then, iteration of the Yule-Walker equations (12) leads to derive the autocovariance values **Γ**_*p*_, **Γ**_*p*+1_ … **Γ**_*q*_ up to the desired lag *q*. Finally, proper rearrangement of the autocovariance values leads to derive the covariances 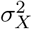, **Σ**_*X*_*q*, and **Σ**_*X*_*q*+1 to be used in (9) and (10) for the computation of the local and global IS.

### 2.4 Non-linear Model-Free Estimation

The k-nearest neighbor (KNN) estimator is a model-free approach which, under the hypothesis of stationarity, defines the local probability density around a given data point as uniform and inversely related to the distance between the point and its *k* nearest neighbors, with *k* that is a parameter controlling the number of neighbors to be calculated [54]. Specifically, given a *d*-dimensional random variable *V* and its *n*-th observation *v*_*n*_ (*n* = 1, …, *N*^*′*^), and denoting as *ϵ*_*n,k*_ twice the distance between *v*_*n*_ and its *k*^*th*^ nearest neighbor in the *d*-dimensional space, the probability mass of the ball of radius *ϵ*_*n,k*_*/*2 surrounding *v*_*n*_ is supposed to be constant and equal to:

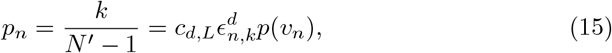

where *c*_*d,L*_ is the volume of the *d*-dimensional unit ball given a norm *L*. From (15), adding a bias-correction term equal to log *k − ψ*(*k*), where *ψ*(*·*) is the digamma function [54], the KNN estimate of the information content of the observation *v*_*n*_ is obtained as:

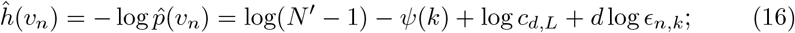

the entropy of *V* is then estimated as the ensemble average [20]:

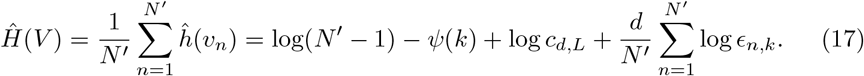

In the computation of the information storage, a naïve estimator would be that applying (16) and (17) to the random variables *x*_*n*_, 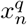, and 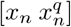 to estimate their information content, and then summing the estimates as in (3) to get the local IS, from which the global IS is derived as in (4). However, this may not be adequate in practice, since the dimension of the analyzed variables is largely different, and the bias of the estimator (16) varies with the dimension. To circumvent this problem, we consider the solution proposed by Kraskov [55] to perform a neighbor search in the highest-dimensional space (here, that spanned by the realizations of 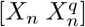) and then using the computed distances in the lower dimensional spaces (here, those spanned by the realizations of *X*_*n*_ and 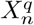) to perform separate range searches. In our context, we compute first the information content of 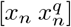 via a simple adaptation of (16):

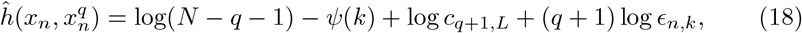

where *ϵ*_*n,k*_ is twice the distance from 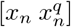 to its *k*^*th*^ nearest neighbor. Then, given the distances *ϵ*_*n,k*_, the information contents in the lower-dimensional spaces are estimated via range searches:

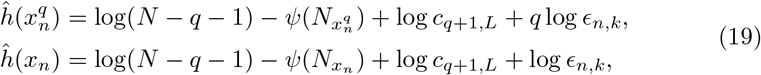

where 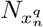 and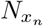 are the number of realizations of 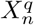 and *X*_*n*_ whose distance from *x*_*n*_ and 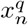 is smaller than *ϵ*_*n,k*_*/*2. Finally, the local IS is computed as 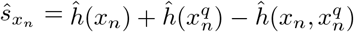, which yields:

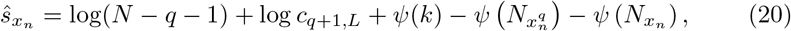

and the global IS results combining (10) and (20):

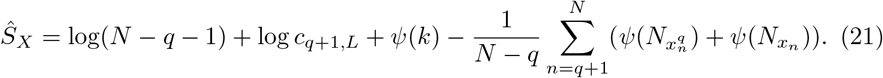

The KNN estimator makes use of only one free parameter, i.e., the number of neighbors *k*, which allows to control the bias-variance tradeoff (larger *k* reduces the variability of estimates, at the expense of a larger bias).

### 2.5 Dataset description and pre-processing

The local measures of regularity described in Sections 2.1-2.4 were applied to EEG signals to assess how the predictability of the brain signals varies in relation to the simultaneously recorded cardiac activity, and therefore to investigate how these two processes overlap in generating physiological events such as the cortical processing of the heartbeat. The aim is to obtain information on the interaction between two physiological signals by analyzing directly only one process (in this case, the EEG reflecting brain activity) and then observing how its properties change when the descriptive index (in this case, the local IS) is evaluated synchronously with the second process (in this case, the one reflecting the heartbeat times).

Signals were acquired at the “Institute for Advanced Biomedical Technologies – ITAB” of the “G. d’Annunzio” University of Chieti-Pescara. Details regarding the dataset can be found in *Zaccaro et al*. [56]. The dataset consists of EEG and ECG signals acquired synchronously on twenty subjects (14 females, age 28.28 *±* 4.78 years), neurologically healthy and not undergoing any psychopharmacological therapy or prolonged drugs assumption. A gel-based BrainAmp EEG system (BrainCap MR, Brain Vision, LLC) with 64 electrodes placed according to the International extended 10/20 system was employed with regard to EEG acquisition (taking the midfrontal electrode as the reference and the inion one as ground) [56], while a one-lead ECG system (BIOPAC Systems, Inc) was used, both with a sampling frequency of 2 kHz. During the acquisition, subjects were in a resting-state condition and were asked to stay seated with eyes open. The study was approved by the Institutional Review Board of Psychology, Department of Psychological, Health and Territorial Sciences, “G. d’Annunzio” University of Chieti-Pescara (Protocol Number 44 26 07 2021 21016), in compliance with the Italian Association of Psychology and the Declaration of Helsinki guidelines and its later amendments. Each subject signed a written informed consent.

EEG signals were pre-processed offline with MATLAB R2021b (The Mathworks, Inc.) using the open source EEGLAB signal processing Toolbox [57]. Signals were bandpass filtered with a Hamming window FIR filter with cutoff frequencies of 0.5 *−* 40 Hz. Artifacts and noise due to movements or incorrect contact of electrodes with skin were manually removed, and the signals from the noisy channels were spherically interpolated. Independent Component Analysis (ICA) was employed (fastICA algorithm [58]) to limit the influence of artifacts on the EEG signals. Finally, signals were subsampled to 128 Hz, in order to reduce the redundancy between consecutive samples prior to the information-theoretic analysis, and re-referenced to the average of all channels [56]. As regards the ECG signals, a modified version of the Pan-Tompkins algorithm [59] was applied to ECG traces of each subject to identify R-peaks; subsequently, T and P waves were extracted using an appositely developed threshold-based peak detection algorithm.

For the subsequent analyses, only 18 subjects were used, due to the presence of artifacts; the length of the recordings was 423.47 ± 27.5 s (range: 320.26 s – 462.29 s). A summary of the pre-processing procedure is reported in the first box of Fig. 1(a).

**Figure 1:**
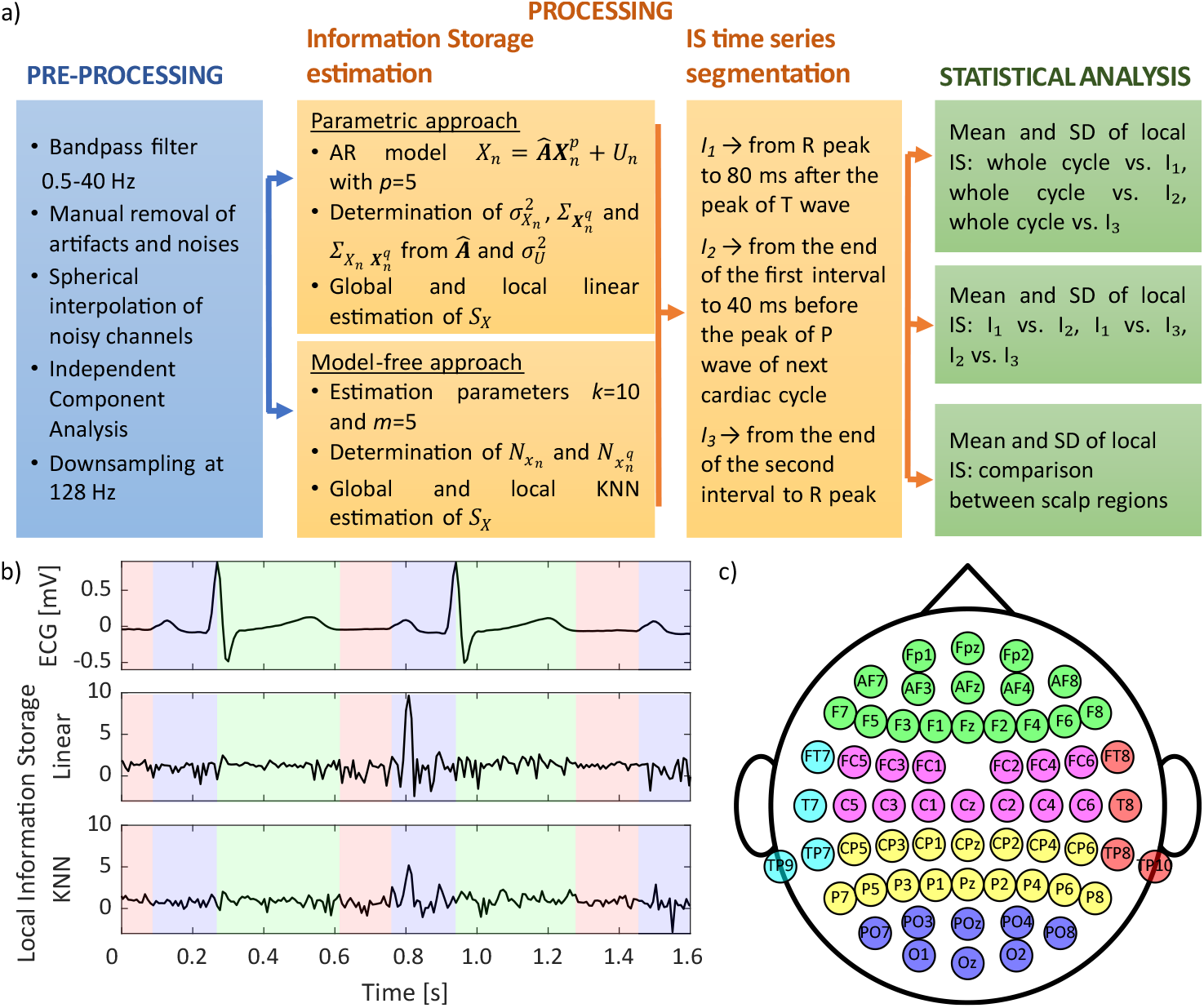
Schematic representation of the methodological approach followed to analyze heartbeat-related Information Storage dynamics. (a) Pipeline of preprocessing (blue box), processing (orange boxes) and statistical analysis (green boxes) steps performed on the EEG signals. (b) Representative portion of the ECG signal and corresponding local information storage time series obtained by linear and KNN estimators, depicting the three time intervals obtained through segmentation of the cardiac cycle (I_1_ in green, I_2_ in red and I_3_ in purple). (c) Schematic of the placement of the EEG electrodes, evidencing the clustering of electrodes according to six scalp regions: frontal (F) in green, central (C) in magenta, parietal (P) in yellow, occipital (O) in blue, right-temporal (Tr) in red and left-temporal (Tl) in cyan.

### 2.6 Data analysis

For each subject, the global and local IS were computed on EEG channels using both the parametric and the model-free approach described in Sections 2.1-2.4 (second box of Fig. 1(a)).

With regard to linear estimation, the AR model was identified for each EEG time series through the well-known least-squares estimator, fixing the model order *p* and the lag *q*. The Akaike (AIC) and Bayesian (BIC) Information Criteria were used to guide the selection of the order *p* of the AR model [60, 61]; however, a minimum in the two figures of merit was not reached, as it often happens when dealing with EEG signals [62]; therefore an order *p* = 5 was selected as the value for which (on average of all subjects and electrodes) the decrement of both AIC and BIC figures of merit moving from an order to the subsequent one was less than 10%. Although the AR model describes interactions between samples up to a lag *p*, typically the correlation function decades to zero with a time lag greater than *p*. Therefore, to account for the whole correlation structure, interactions between the present and past samples were accounted up to a lag *q* such that the autocorrelation vanishes [27]; in this work, *q* = 10 was chosen to ensure a spectral radius of the AR process smaller than 10^*−*8^ [27]. After AR model identification, the variance and the covariance matrices needed to compute local and global IS were obtained as described in Section 2.3.

With regard to the model-free estimator, the number of neighbors *k* was set according to previous studies dealing with short-time series [20, 63], i.e., *k* = 10. The dimension *q* of the embedding vector representative of the process history was set in accordance to the AR model order parameter *p* determined for the linear estimator, taking into account that higher values are not recommended in model-free analyses in order to limit the curse of dimensionality [29]. In our analysis, we used the Chebyshev norm taking the maximum distance between the scalar components of the realizations of *V*, so that the volume *c*_*d,L*_ is equal to 1 and the term log *c*_*q*+1,*L*_ vanishes in (20) and (21).

For each EEG time series, the local IS was analyzed separately in three different intervals determined by segmenting each detected cardiac cycle as depicted in the third box of Fig. 1(a) and in Fig. 1(b). Segmentation was performed to take into account the influence of the Cardiac Field Artifact (CFA) on brain signals. As reported in previous works [64], the cardiac electrical activity can be detected over the entire body surface, including the scalp, and leads to changes in the amplitude of EEG signals. The influence of the CFA on the EEG varies with the spatial distribution on the scalp, depending on the proximity of the electrodes to the heart, but also with time, depending on the corresponding phase of cardiac cycle [64]. These remarks have been considered for defining the three intervals. Specifically, the first interval (I_1_) starts at the R-peak of the ECG signal and ends 80 ms after the peak of T-wave, the second one (I_2_) starts at the end of the first interval and ends 40 ms before the peak of the P-wave of the next cardiac cycle and, finally, the third interval (I_3_) corresponds to the remaining part of the cardiac cycle until the R-peak of the subsequent cycle. Given these temporal windows, the second interval I_2_ may be considered as low-CFA segment; in other words, the influence of CFA on the EEG signals can be considered almost negligible within this interval. On the other hand, the other two intervals, I_1_ in particular, are most affected by the artifact given that the cardiac electric field is more evident during the QRS complex and the T-wave [64]. In order to gain consistency, we took the first 300 heartbeats into account for the analysis, also in accordance to the standards of short-term heart rate variability analysis [35]. For each subject and electrode, the mean and the standard deviation (SD) of the local IS measures were computed within each of the three intervals, thus obtaining 300 values of local IS mean and standard deviation. To provide a reference unrelated to time segmentation, the mean and SD of the local IS were computed also within the whole cardiac cycle.

### 2.7 Statistical analysis

The statistical analysis was performed considering the mean and SD of the local IS computed within each whole cardiac cycle or within one of the three identified intervals, using either the linear parametric estimator or the non-linear model-free estimator (see Fig. 1). A single value of the mean local IS and of the SD of the local IS was obtained for each analyzed EEG signal taking the average over the 300 cardiac cycles considered. Then, the distributions across subjects of the two measures were compared using the parametric Student’s t-test as follows. For all the analyses, the significance level was set to *p* = 0.05.

The first analysis was aimed at comparing, for the EEG signal measured from a given electrode, the distributions across subjects of the mean local IS and of the SD of the local IS obtained for the whole cardiac cycle versus each of the distributions computed within the three intervals, i.e., comparing G vs I_1_, G vs I_2_ and G vs I_3_. In this case, the paired Student’s t-test was followed by Bonferroni correction, with *n* = 3 comparisons.

The second analysis was aimed at comparing, for the EEG signal measured from a given electrode, the distributions across subjects of the mean local IS and of the SD of the local IS obtained in the three intervals, i.e., comparing I_2_ vs I_1_, I_3_ vs I_1_ and I_3_ vs I_2_. Also in this case, the paired Student’s t-test was followed by Bonferroni correction, with *n* = 3 comparisons.

Moreover, to analyze spatial variations across the scalp, we identified six regions according to the electrode locations reported in Fig. 1(c), i.e., frontal (F), central (C), parietal (P), occipital (O), right-temporal (Tr) and left-temporal (Tl); for each subject and each interval, values of the mean local IS and of the SD of the local IS were determined for each region taking the average of the values referred to all the electrodes belonging to that region. The comparison between pairs of regions was performed using the paired Student’s t-test with Bonferroni correction for multiple comparisons (*n* = 15).

Finally, a measure of the effect size was also determined to assess the magnitude of the differences observed among regions. Specifically, denoting with 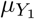 and 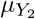 and with 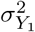 and 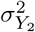 the mean and the variance of two distributions *Y*_1_ and *Y*_2_ obtained measuring the mean local IS or the SD of the local IS across subjects, we computed the Cohen’s *d* measure defined as follows [65]:

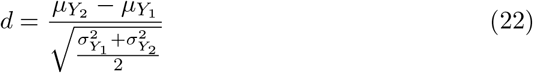

onventionally a small effect size occurs for *d* = 0.2, a medium effect size for *d* = 0.5 and a large effect size for *d* = 0.8 [65].

## 3 Results

Figure 2(a) depicts the scalp distribution of the mean values of the global IS (G, computed over the whole cardiac cycle) and the local IS (computed within each interval I_1_, I_2_, or I_3_) obtained using the linear estimator. The scalp distribution of the mean values is very similar comparing the global and local IS, and does not depend on the considered interval. The regularity reflected by the mean local IS is higher at the parietal and occipital regions if compared with temporal regions. These observations are supported by the statistical analysis comparing the distribution of the mean values of the global IS with each of the distributions of the local IS, or comparing pairs of distributions of the local IS, depicted respectively in Fig. 2(b) and in Fig. 2(c). Indeed, slight statistically significant differences (*p* = 0.0084 *±* 0.0041) were found for only a few electrodes in the frontal, occipital and right temporal regions.

**Figure 2:**
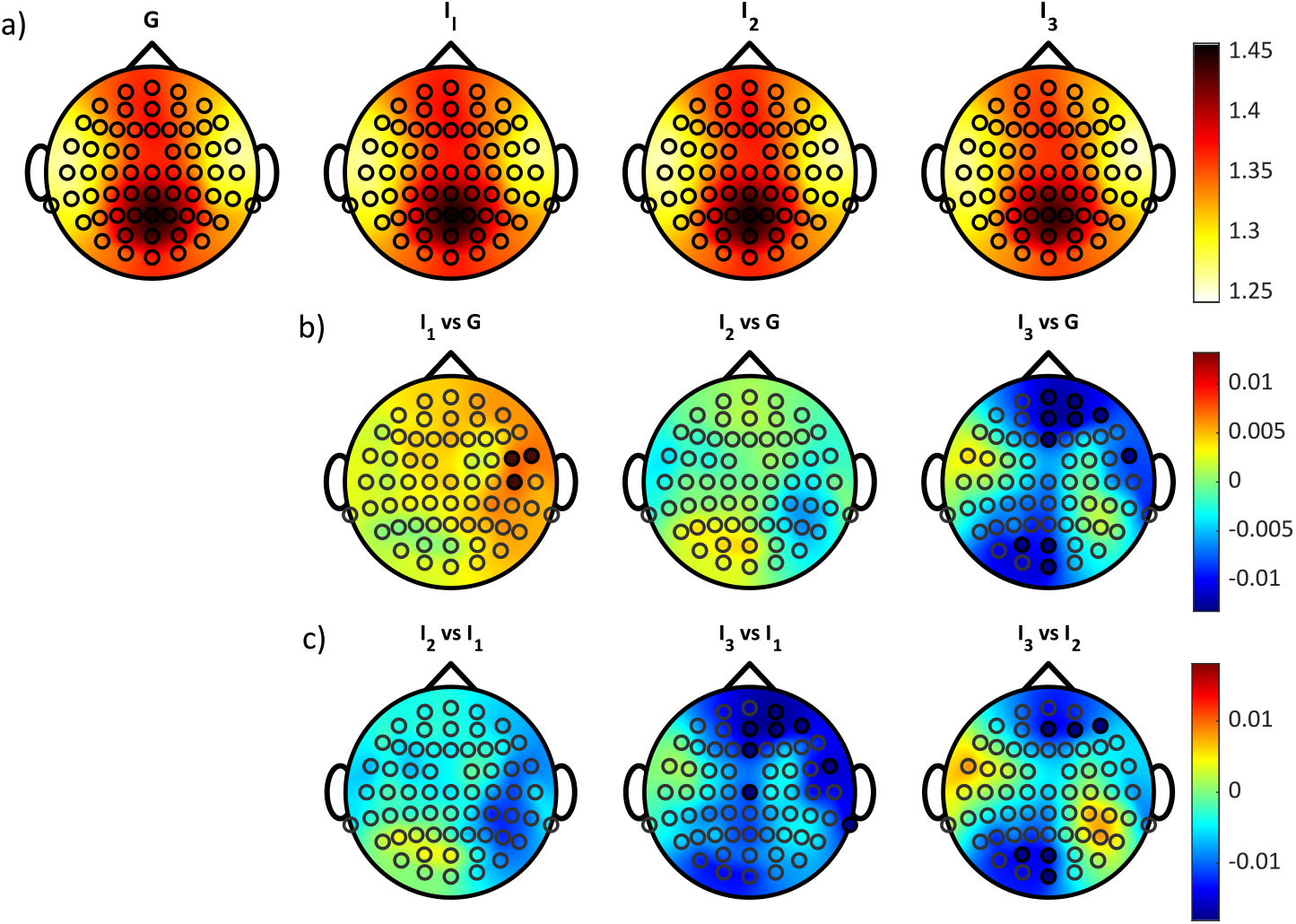
Analysis of the mean of the local IS of the EEG computed using the linear estimator. (a) Distribution over the scalp of the average values across subjects of the mean of the global IS (G, computed over the whole cardiac cycle) and of the mean of the local IS (computed in a specific interval I_1_, I_2_, or I_3_). (b) Distribution over the scalp of the average values across subjects of the difference between the mean global IS and the mean local IS computed within each interval I_1_, I_2_, I_3_. (c) Distribution over the scalp of the average values across subjects of the difference between the mean local IS computed within pairs of intervals I_1_, I_2_, I_3_. Red (blue) electrodes show positive (negative) differences (paired Student’s t-test with Bonferroni correction, *p <* 0.05*/n*, with *n* = 3 comparisons). No statistically significant differences were found.

Figure 3(a) reports the scalp distribution of the SD values of the global and local IS measures computed through the linear estimator. The variability of the local IS exhibits a different pattern if compared to the mean, with SD values more uniformly distributed over the scalp and more dependent on the temporal windows over which they are computed. As shown in Fig. 3(a), the variability of the global measure obtained computing the SD of the local IS over the whole cardiac cycle is markedly higher than the variability of the local measure computed over the three intervals. This trend can be evidenced from Fig. 3(b), showing that the difference between the SD of the global and local IS values is always lower than zero and is statistically significant at each location in the scalp. Moreover, the variability of the local IS shows a decreasing trend going from the first to the third interval, as documented in Fig. 3(c), where statistically significant differences are observed comparing I_3_ vs I_2_ and even more comparing of I_3_ vs I_1_; the differences between the first two intervals (I_2_ vs I_1_) are statistically significant only for one electrode, being the resulting *p*-value furthermore close to the significance threshold (*p ∼* 0.0064).

**Figure 3:**
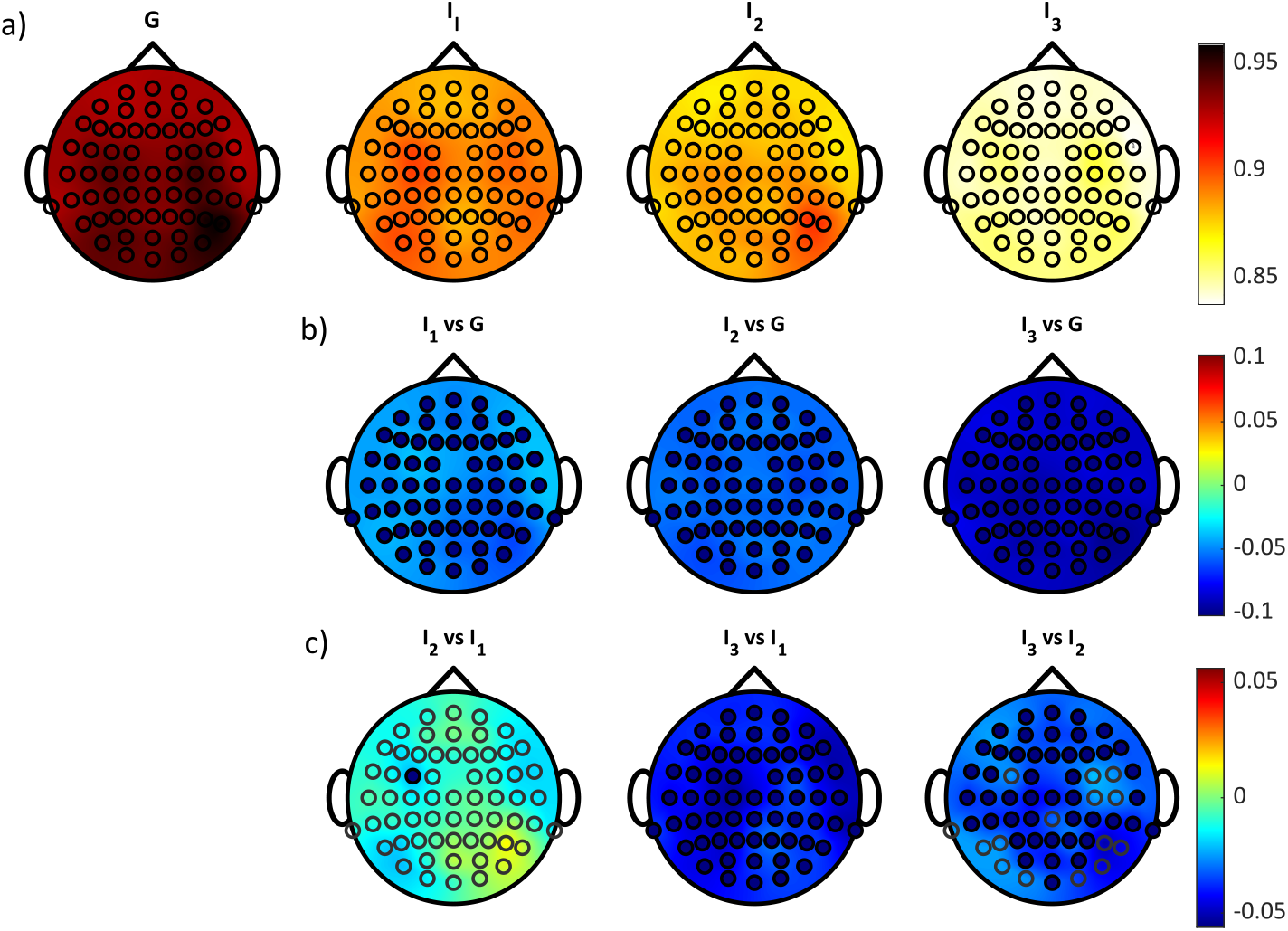
Analysis of the standard deviation (SD) of the local IS of the EEG computed using the linear estimator. (a) Distribution over the scalp of the average values across subjects of the SD of the global IS (G, computed over the whole cardiac cycle) and of the SD of the local IS (computed in a specific interval I_1_, I_2_, or I_3_). (b) Distribution over the scalp of the average values across subjects of the difference between the SD of the global IS and the SD of the local IS computed within each interval I_1_, I_2_, I_3_. (c) Distribution over the scalp of the average values across subjects of the difference between the SD of the local IS computed within pairs of intervals I_1_, I_2_, I_3_. Red (blue) electrodes show statistically significant positive (negative) differences (paired Student’s t-test with Bonferroni correction, *p <* 0.05*/n*, with *n* = 3 comparisons).

Figs. 4 and 5 report the scalp distribution of the mean and SD of the global and local IS computed using the model-free estimator implemented with the nearest neighbor technique. Results evidence that the nonlinear estimator leads to values of the local information storage and of its variability lower than those obtained using the linear approach (cf. Fig. 4 vs Fig. 2 and Fig. 5 vs Fig. 3). Nevertheless, the spatial distribution and the changes across conditions displayed by the mean and SD of the local IS computed with the model-free estimator are very similar to those obtained for the parametric estimator. Indeed, the mean IS has very similar scalp distribution when computed globally over the whole cardiac cycle or locally within the three intervals of the cycle (Fig. 4(a)), and only few statistically significant differences are found in the comparisons of global and local measures as well as in that of local measures in the three intervals (Fig. 4(b,c)) with *p*-values close to the threshold (*p* = 0.0087 *±* 0.0062). The SD of the local IS shows values more uniformly distributed across the scalp (Fig. 5(a)), significantly higher when computed over the whole cardiac cycle than over the intervals (Fig. 5(b)), which tend to decrease moving from the first to the second and third interval. Specifically, the differences between the intervals were found to be significant over the whole scalp when comparing I_3_ and I_1_, and mostly also when comparing I_3_ and I_2_, while in the comparison between I_2_ and I_1_ the significant differences are mainly related to the temporal regions (Fig. 5(c)).

**Figure 4:**
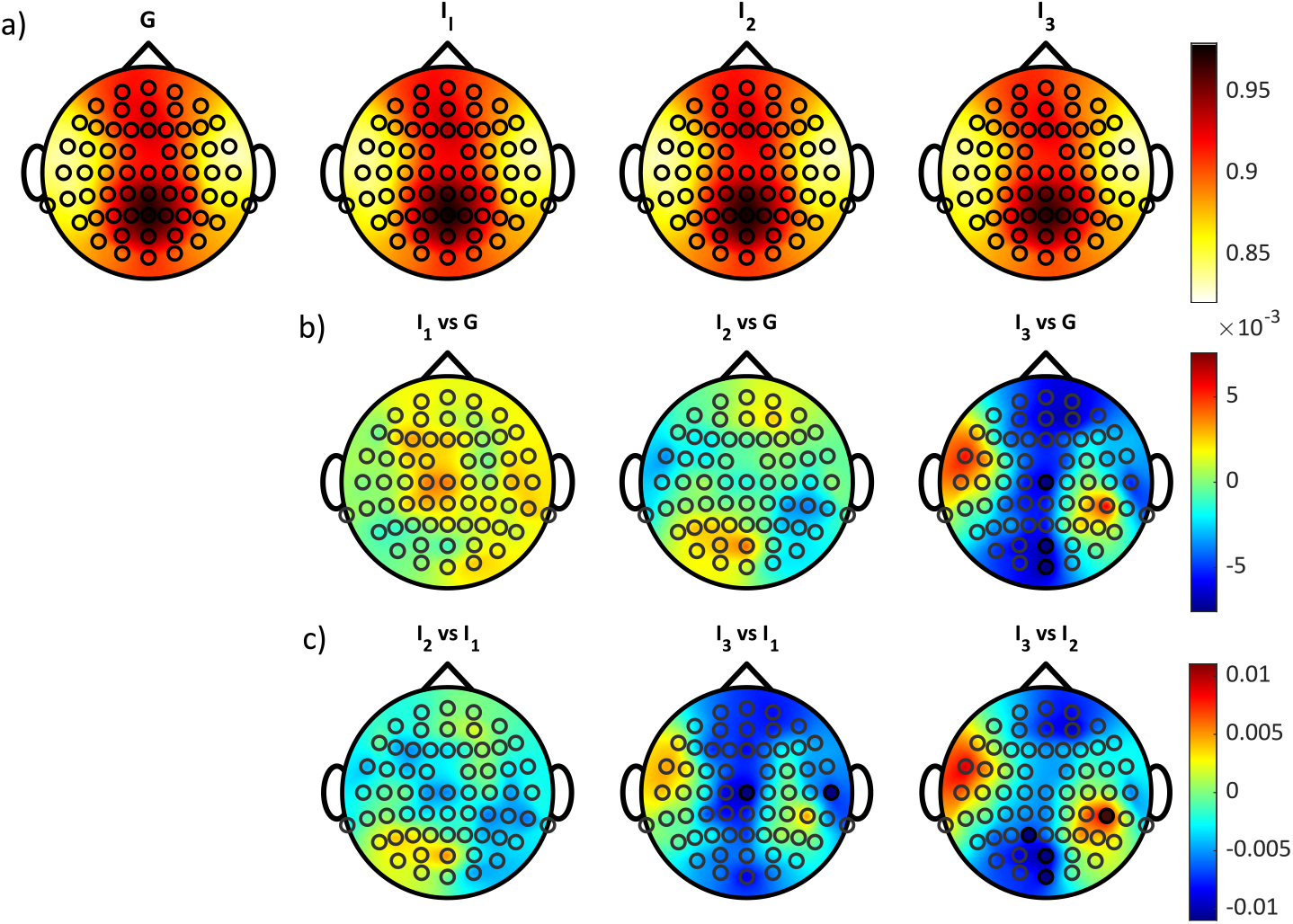
Analysis of the mean of the local IS of the EEG computed using the k-nearest neighbor estimator. (a) Distribution over the scalp of the average values across subjects of the mean of the global IS (G, computed over the whole cardiac cycle) and of the mean of the local IS (computed in a specific interval I_1_, I_2_, or I_3_). (b) Distribution over the scalp of the average values across subjects of the difference between the mean global IS and the mean local IS computed within each interval I_1_, I_2_, I_3_. (c) Distribution over the scalp of the average values across subjects of the difference between the mean local IS computed within pairs of intervals I_1_, I_2_, I_3_. Red (blue) electrodes show positive (negative) differences (paired Student’s t-test with Bonferroni correction, *p <* 0.05*/n*, with *n* = 3 comparisons). No statistically significant differences were found.

**Figure 5:**
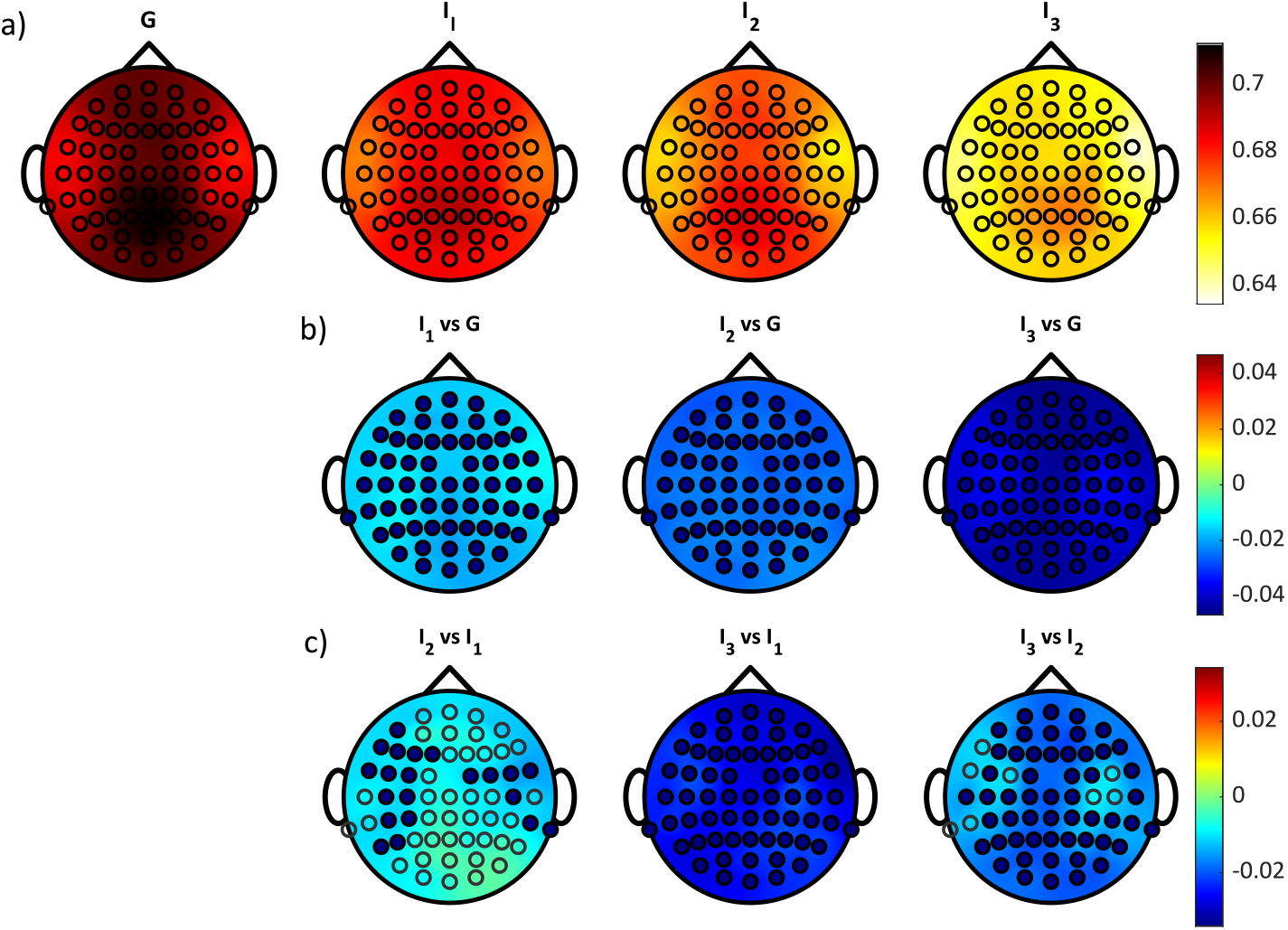
Analysis of the standard deviation (SD) of the local IS of the EEG computed using the k-nearest neighbor estimator. (a) Distribution over the scalp of the average values across subjects of the SD of the global IS (G, computed over the whole cardiac cycle) and of the SD of the local IS (computed in a specific interval I_1_, I_2_, or I_3_). (b) Distribution over the scalp of the average values across subjects of the difference between the SD of the global IS and the SD of the local IS computed within each interval I_1_, I_2_, I_3_. (c) Distribution over the scalp of the average values across subjects of the difference between the SD of the local IS computed within pairs of intervals I_1_, I_2_, I_3_. Red (blue) electrodes show statistically significant positive (negative) differences (paired Student’s t-test with Bonferroni correction, *p <* 0.05*/n*, with *n* = 3 comparisons).

Fig. 6 shows the results of the analysis carried out on the mean values of the local IS computed in each of the three analyzed intervals using both linear (first row) and KNN (second row) estimators and averaged over the electrodes of the six detected regions of the scalp. Results evidence that, for both estimators, the local IS computed in the first two intervals is significantly lower in the temporal regions than in the other areas; a similar pattern is observed for the mean local IS computed during I_3_, though with a lower number of statistically significant differences. The effect size measure highlighted that the most relevant differences are between the temporal region and the parietal/occipital regions: considering the average values among the three intervals, when comparing the temporal region with the parietal and occipital, the Cohen’s *d* is respectively 1.05 and 1.02 using the linear estimator, and 0.83 and 0.78 using the KNN estimator.

**Figure 6:**
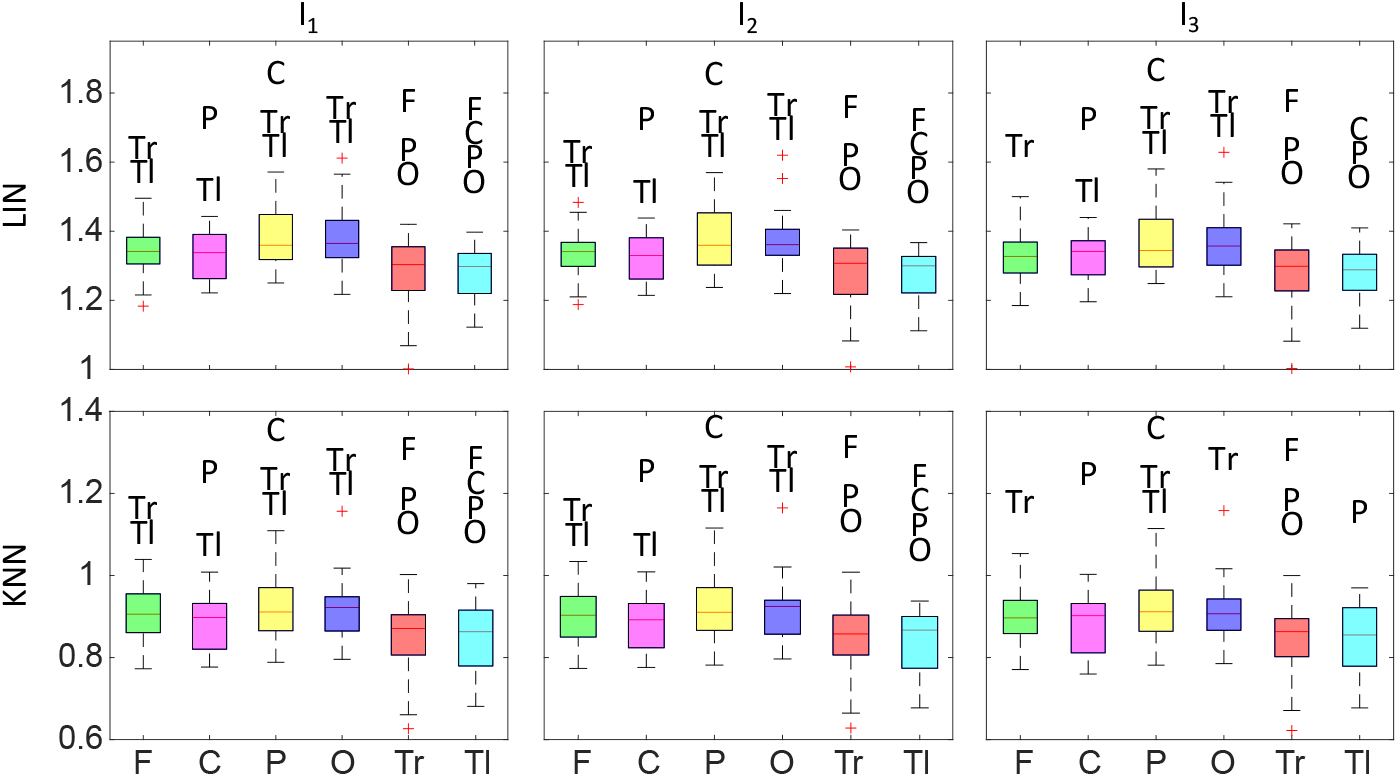
Boxplots of the mean values of the local IS computed in each of the three intervals (I_1_, I_2_, I_3_) using the linear (LIN) and k-nearest neighbor (KNN) estimator for the six scalp regions: frontal (F), central (C), parietal (P), occipital (O), temporal right (Tr) and temporal left (Tl). Statistical analysis: region name, *p <* 0.05*/n*, paired t-test with Bonferroni correction (*n* = 15 comparisons).

Fig. 7 depicts the results of the analysis of the regional differences carried out using the standard deviation of the local IS. During I_2_, the linear estimator shows significantly higher variability of the local regularity in the parietal region when compared to frontal and right temporal regions. In this case, a higher number of statistically significant differences is detected using non-linear estimator; for all the three intervals, the variability in the parietal region is significantly higher than in the left temporal area.

**Figure 7:**
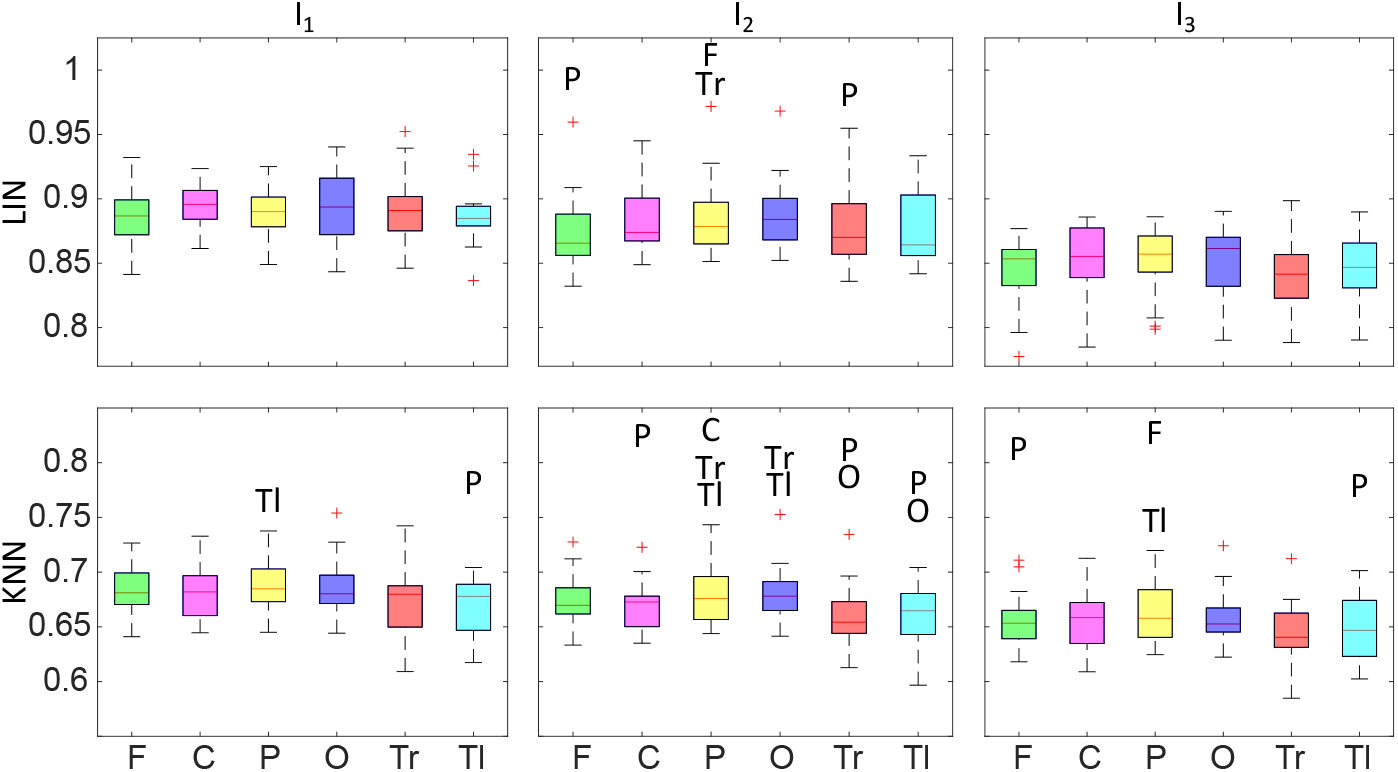
Boxplots of the standard deviation of the local IS computed in each of the three intervals (I_1_, I_2_, I_3_) using the linear (LIN) and k-nearest neighbor (KNN) estimator for the six scalp regions: frontal (F), central (C), parietal (P), occipital (O), temporal right (Tr) and temporal left (Tl). Statistical analysis: region name, *p <* 0.05*/n*, paired t-test with Bonferroni correction (*n* = 15 comparisons).

## 4 Discussion

This study proposes a novel approach for investigating the temporal evolution of the complexity of biosignals measured from physiological systems. The use of measures able to reveal the patterns of the information transferred, stored and modified within dynamical systems in a time-resolved way [22, 30, 31] allows a more thorough assessment of the physiological control mechanisms of individual organs as well as their interactions. In this work, the peculiarity of local information measures was exploited to look at the variability of the regularity of neural signals related to different phases of the cardiac cycle, providing an alternative method to study brain-heart interactions. Our results regarding the global IS and the mean values of the local IS of the EEG identify the scalp areas showing the more regular neural activity during a resting state condition, although they are unable to discriminate the time window with the strongest influence of the cardiac activity on the EEG regularity. On the other hand, the standard deviation values of the local IS are almost uniformly distributed over the scalp but allow discriminating different behaviors within heartbeats, being thus more useful to reflect cortical processing of the cardiac cycle.

### 4.1 Mean response

The scalp distribution of the mean values of the local information storage shown in Figs. 2(a) and 4(a) may be related to the continuous communication between heart and brain during a resting-state condition, which is equally reflected in the global IS measure. In fact, higher values of the information stored in EEG signals, indicative of a more regular neural activity, could suggest the presence of a prevalent neural rhythm even in the absence of tasks, as already demonstrated in a previous work through simultaneous acquisition of EEG and fMRI signals [49]. A correlation between neural activity in regions of the DMN and the spectral power of EEG signal has been previously observed documenting that an increased activity in vmPFC, PCC and PLC is associated with an augmented EEG power in the alpha and beta bands in parietal and occipital regions [49]. Similar considerations can be made from our results reported in Fig. 6, since the average values of the local IS in the temporal regions are significantly lower than in parietal and occipital areas. Such results are similar to those reported in [66], showing how the neural response to the heartbeat is restricted to the parietal regions of the scalp.

The similarity of the results obtained with global and local measures may suggest that the functional mechanisms determining the regularity of neural rhythms are the same throughout the whole cardiac cycle. Indeed, since the intervals were defined by evaluating the different influence of the CFA at each heartbeat and knowing that the influence of this artifact is prevalent in the first and third interval [64], the similar IS average values detected in all the time windows suggest that the cardiac activity does not have an influence on brain regularity.

### 4.2 Standard deviation response

Our results show that the parameter most influenced by the course of time is the variability of the information stored in the EEG signals, as demonstrated by the average values of the IS standard deviation reported in Figs. 3(a) and 5(a). Indeed, both the linear estimator and the nearest neighbor approach show how the standard deviation of IS changes markedly if evaluated globally or locally within the time windows corresponding to the different phases of the cardiac cycle. In particular, there is a significant decrease of this quantity when going from the first to the third interval, hence suggesting that the variability in the EEG regularity is strongly affected for each heartbeat by the cardiac pulse occurring close to the R-peak, which acts as a trigger. The variability of the information stored in the EEG signals then decreases with time, thus suggesting that the effect of the stimulus (heartbeat) diminishes and is related to a reduced perturbation of the IS measure till the occurrence of the successive stimulus. These results highlight that the variability of the EEG regularity is pivotal in the study of the mechanisms of brain-heart communication. This finding reinforces the feasibility of employing the variability of local information measures to differentiate the behavior of interacting systems in the transition across different states, as already been demonstrated in [33] on cardiac and respiratory signals using the local Granger Causality.

### 4.3 Comparison between linear and non-linear estimations

The local and global measures of EEG regularity were computed using both parametric and model-free approaches, implemented through the linear and the k-nearest neighbors estimators, respectively. The estimated values can depend on the method employed, given that a model-based parametric estimator cannot suitably capture nonlinearities and should be in principle limited to assess linear dynamics, contrary to the model-free approach which is more robust to the presence of non-stationarities, but in turn is less computationally reliable [1].

The results obtained through parametric and model-free approaches, both in terms of mean (Figs. 2 and 4) and standard deviation (Figs. 3 and 5), evidenced similar spatial and temporal distributions of the local IS. However, the values obtained by the nearest neighbor estimator are generally lower than those obtained using the linear estimator. The reasons of such a difference are difficult to explain and may be related to several aspects. A possible explanation may be the presence of local non-linearities and/or non-stationarities, which have been shown determine a temporal variation of local information measures computed using globally identified linear regression models in the case of bivariate causality measures [33], which could play a role also in an univariate setting like in our case; even if a visual inspection of EEG signals has been carried out before performing the analysis in order to check global stationarity, local non-stationarities may still be present in EEG signals, as evidenced by previous studies in the literature [67]. Moreover, it is not possible to exclude the effect of a bias in the nonlinear measures obtained using the KNN estimator, due to the difficulty of working on high-dimensional spaces (the embedding dimension was set at 5). Nevertheless, in spite of the different absolute values, the well-comparable changes across windows obtained with the two estimators both for the mean and the variability of the local IS suggest that non-linearities do not play a crucial role in determining the EEG regularity in relation to the heartbeat stimuli. In view of this, we conclude that a parametric approach appears to be suitable enough to characterize the patterns of local IS in various scalp regions and across different phases of the cardiac cycle.

Another issue to take into account when discussing the results is the proper choice of the parameters of the estimators. We underline that in our analysis an optimal order of the AR model could not be found, suggesting that this class of models may be unsuitable to fully describe the EEG dynamics. It has been indeed observed that the correct AR model order to describe EEG signals would ideally be infinite [62], and that the class of Autoregressive-Moving Average (ARMA) models that also takes into account autocorrelations of errors should be used instead [68]. Here, we stick to the AR representation because of the difficulty to identify ARMA models and to provide the local formulation of information measures for this class of models. Nevertheless, the parametric representation and the exploitation of the extended Yule-Walker equations allow to take into account long lags in the representation of the past history of the process, covering long memories until the covariance decays, while this is not possible using the model-free approach [69]. In fact, the estimation of information measures is more difficult using the nearest neighbor estimator and should thus be limited to a small number of past lags; the obtained results in this case could be strongly affected by the choice of the dimension of the embedding vector as well as by the number of neighbors.

### 4.4 Comparison with HEP

Several and somewhat conflicting hypotheses have been reported to describe the complex communication mechanisms between the heart and the brain, being the most supported one that this process is generated by the superposition of multiple events that come into play in different ways and at multiple times [40]. The continuous processing of stimuli from within the body has been also shown to crucially shape our first-person experience [34, 40], allowing to coordinate and unify the perception of the external stimuli to which, even unconsciously, everybody is constantly exposed. This idea emerged from EEG studies focusing on the cortical processing of cardiac signals through measures of the Heartbeat Evoked Potential (HEP) [70, 71, 72]. Depolarization and repolarization of the heart constitutes a visceral stimulus for the brain which is processed in cortical areas like any other stimulus that reaches the body from the external environment. The cortical response to heartbeat is generally investigated as those evoked by any external stimulus, which are known as Event-Related Potentials. Specifically, EEG traces are segmented and timed with respect to the ECG R-peak and then averaged to obtain the waveform of the relative potential [40, 73]. Several studies adopting the HEP led often to contrasting results depending on the task carried out by the subjects under examination [74].

The methodology of analyzing HEPs is markedly different from the approach followed in this paper. Indeed, while our methodology allows to find out the presence of repetitive EEG patters related to the heartbeat, using the HEP any reference to local regularity patterns is instead superseded by the averaged trend of the conduction of the cardiac electrical stimulus on the scalp. For this reason, a direct comparison of our results with other studies that used the HEP is difficult to be carried out, since they operate in a “reverse” way. Nevertheless, there is a quite good agreement between the interval in which we assume there is less influence of merely electric field effects and thus more neural, i.e., I_2_, and the interval in which the HEP is more commonly reported. In fact, although the HEP potential curve strongly depends on data, a large number of studies employing non-parametric cluster-based permutation techniques restricted the influence of heartbeats on neural activity at the fronto-central electrodes and in a time interval between 300 and 600 ms after the ECG R-peak [40, 74], which is mostly overlapping with the second time window employed in our analysis, although it is not completely equivalent. A thorough comparison of the two approaches, highlighting similarities and differences, will be the scope of future studies aiming to provide a comprehensive analysis of heartbeat-induced cortical dynamics.

### 4.5 Limitations and future perspectives

The approach proposed in this work overcomes the limitations of the commonly employed global information measures which cannot provide point-in-time insights about physiological dynamics. Despite this, the method applied in the present study for investigating cortical processing of heartbeat has some limitations which should be taken into account.

A first limitation derives from the employment of ICA for data preprocessing. The ICA approach is very effective for separating the components associated with neural and non-neural sources from EEG signal, allowing to filter out non-neural information. Despite using ICA is a well-established practice in HEP analysis [74], its application may lead to remove not only the CFA, but also useful information related to the cortical processing of the heartbeat [40] being both generated by the same source. To overcome such limitation, an alternative methodological approach that could be used is the Current-Source Density (CSD) transformation, which allows to reduce the presence of the artifacts from EEG signals without being so limiting with regard to the analysis of the HEP and therefore to the cortical processing of the heartbeat [75, 76]. Other studies instead remove the CFA by using a bipolar reference, given that subtracting the signal from adjacent electrodes corresponds to eliminate any signal coming from common external or non-neural sources [46].

Another limitation concerns the assumption of global stationarity that is made when using the parametric and non-parametric approaches developed in this work. In this context, future developments could be aimed to compare the results achieved with the local analysis carried out in this paper with a time-varying approach [77] that overcomes the assumption of signal stationarity, while also allowing for high time resolution information as the local analysis [30]. In addition, the analysis of variations of neural activity regularity could be also carried out not on the EEG scalp signals, but instead after performing the reconstruction of the sources [28]. Another analysis envisaged is the application of the approach developed in this work to data acquired during the execution of tasks generally used for studying the HEP (e.g., the heartbeat counting task [40, 71]), or in presence of pathological conditions (e.g., depression, nightmare disorder [40, 78, 79]), in order to allow not only to identify differences among conditions, but also to compare the results to those already reported in the literature.

## 5 Conclusions

In this work, the application of a local measure of information storage has allowed to assess brain-heart interactions – i.e., interactions occurring in a bivariate process – starting from the univariate analysis of a signal with faster dynamics (i.e., the EEG) and its timing with a slower one (i.e., the ECG). From a physiological point of view, our results have demonstrated that the local information storage of EEG signals allows to assess the variation of brain rhythms regularity within the cardiac cycle, detecting mechanisms of heart-brain interactions potentially complementary to those typically studied through HEP analysis. Our findings have evidenced that the heartbeat is capable of evoking alterations in the information processed by the electric brain activity even during resting-state conditions, manifested mainly through changes in the standard deviation of the local information storage in the EEG signal rather than in its mean values.

Therefore, local information measures – still not widely employed given their recent development – could represent an important tool for the analysis of physiological mechanisms, since the localization in time of the information carried and processed within a physiological signal could be exploited to identify internal dynamics of the human organism. This may contribute to shed more light to the physiological mechanisms underpinning the cortical processing of the heartbeat, still highly debated in the literature.

## CRediT authorship contribution statement

**Chiara Barà**: Software, Formal analysis, Investigation, Writing - Original Draft, Visualization. **Andrea Zaccaro**: Data Curation, Writing - Review & Editing. **Yuri Antonacci**: Validation, Formal analysis, Writing - Review & Editing. **Matteo Dalla Riva**: Formal analysis, Writing - Review & Editing. **Alessandro Busacca**: Resources, Writing - Review & Editing. **Francesca Ferri**: Resources, Data Curation, Writing - Review & Editing. **Luca Faes**: Conceptualization, Methodology, Software, Writing - Review & Editing, Supervision. **Riccardo Pernice**: Methodology, Software, Formal analysis, Writing - Review & Editing, Supervision.

